# Intracranial Electric Field Recording During Multichannel Transcranial Electrical Stimulation

**DOI:** 10.1101/2021.12.19.473336

**Authors:** Minmin Wang, Jiawei Han, Hongjie Jiang, Junming Zhu, Wuwei Feng, Pratik Y. Chhatbar, Jianmin Zhang, Shaomin Zhang

## Abstract

**Background:** Multichannel transcranial electrical stimulation (tES) modeling and optimization have been widely studied in recent years. Its theoretical bases include quasi-static assumption and linear superposition. However, there is still a lack of direct *in vivo* evidence to validate the simulation model and theoretical assumptions.

**Methods:** We directly measured the multichannel tES-induced voltage changes with implanted stereotactic-electroencephalographic (sEEG) electrodes in 12 epilepsy subjects. By combining these measured data, we investigate the linear superposition and prediction accuracy of simulation models for multi-electrode stimulation and further compare the induced EF differences between transcranial direct current stimulation (tDCS) and transcranial alternating current stimulation (tACS).

**Results:** Our *in vivo* measurements demonstrated that the multi-electrode tES-induced voltages were almost equal to the sum of the voltages generated independently by bipolar stimulation. Both measured voltages and electric fields obtained *in vivo* were highly correlated with the predicted values in our cohort (Voltages: *r* = 0.92, *p* < 0.001; electric fields: *r* = 0.74, *p* < 0.001). Under the same stimulation intensity, the tDCS-induced peak-zero voltages were highly correlated with the values of tACS (*r* = 0.99, *p* < 0.001; *s* = 0.99).

**Conclusions:** The *in vivo* measurements provides confirmatory results for linear superposition and quasi-static assumption within the human brain. Furthermore, we found that the individualized simulation model reliably predicted the multi-electrode tES-induced electric fields.

## 1. Introduction

As an increasingly popular method for non-invasive brain stimulation (NIBS), transcranial electrical stimulation (tES) has been widely used in multiple areas of clinical research, such as psychiatric disorders [1] and stroke rehabilitation [2]. However, the clinical effects of tES are inconsistent [3, 4]. Many studies have found that the tES-induced responses can vary substantially across subjects [5, 6]. The variation may be caused by many factors, such as the stimulation setup, anatomical variations, and brain state. The tES-induced spatial electric fields (EFs) within the corresponding brain regions are considered a significant factor for the biological effects. Some studies have demonstrated the effects of electric fields on neural activity [7–9]. However, it is difficult to obtain electric field distributions directly from live human brains during electrical stimulation.

Therefore, simulation models have been proposed as an alternative to predict spatial EF distribution within the brain [10, 11]. Furthermore, based on the principle of linear superposition adopted by tES modeling, which implies that the total tES-induced EFs can be expressed as a linear combination of the fields created by each bipolar configuration, the stimulation parameters can be optimized to maximize the electric field intensity or focality in a specific target cortical region [12–14]. A novel 4 × 1 high-definition tES (HD-tES) protocol was proposed and widely used because of its higher focality than traditional tES in the simulated EF [15]. However, despite the application of tES modeling and optimization, there are only a few *in vivo* studies to validate the accuracy of these models, especially for multichannel tES. Some previous studies have reported *in vivo* intracranial measurements of EF in humans or nonhuman primates for bipolar transcranial alternating current stimulation (tACS) [16–18]. To the best of our knowledge, only one study has reported intracranial EF measurements in two non-human primates during multichannel tACS [19]. Multichannel tES-induced EF measurements in the live human brain have not yet been reported. Overall, there is still a lack of direct evidence from *in vivo* measurements to validate the linear superposition of the EF induced by multichannel tES.

Meanwhile, the head tissues are considered predominantly resistive at relatively low stimulation frequencies (<1 kHz) in most related tES modeling, so that the induced electric fields can be described in a quasi-static regime [20]. In this case, under the same stimulation intensity, the simulated peak EF was equal for tES at different stimulation frequencies. Some studies have found that the peak voltages induced by equal-intensity tACS showed no significant difference when the stimulation frequency was increased from 0.5 to 200 Hz [17, 21]. But it is still not clear that whether there are differences in the induced peak EF between the same intensity tACS and transcranial direct current stimulation (tDCS). Although some studies have reported the measurement of tDCS-induced voltage using two deep brain stimulation (DBS) electrodes in the human brain [22, 23], there is still a lack of tDCS-induced EF measurements in the human brain with multiple implanted electrodes.

This study directly measures multichannel tES-induced voltage changes using implanted stereo-electroencephalogram (sEEG) electrodes in epilepsy subjects. With these measured data, we investigated the linear superposition and prediction accuracy of simulation models for multi-electrode stimulation, and further compared the difference in the induced EF between tDCS and tACS.

## 2. Methods

### 2.1 Human subjects

Experimental data were obtained from twelve right-handed epilepsy subjects (four females, 30.8 ± 12.3 years [range 19 - 56]) who underwent intracranial recording to identify the localization of the seizure onset zone. Inclusion criteria were: (1) undergoing intracranial recording to localize sites of seizure initiation for epilepsy surgery; (2) cognitive ability to complete questionnaires; (3) ability to provide informed consent. Exclusion criteria were: (1) cognitive impairment (IQ<70); (2) documented severe depression or other neuropsychiatric diseases; (3) scalp/skin disease; (4) frequent electro-clinical seizures within 24 hours immediately after electrode implantation; (5) space-occupying intracranial lesion. All procedures were approved by the Institutional Review Board at the Second Affiliated Hospital, Zhejiang University School of Medicine, and were performed in accordance with the Declaration of Helsinki. A written consent was obtained from all subjects before participation.

### 2.2 Stimulation protocol

A multichannel transcranial current stimulation device (Starstim 8, Neuroelectrics, Barcelona, Spain) was used in our study. To compare the difference between tDCS and tACS, we tested stimulation current of 0.5 and 1 mA, DC and 100 Hz. Each single tES trial lasted less than 2 min, including a ramp-up time of 10 seconds, stimulation time of 60 seconds and ramp-down time of 5 seconds. To assess the stability of tES, the stimulation process was repeated twice using the same stimulation parameters. Circular Ag/AgCl electrodes (electrode size was 3.14 cm2) and conducting gel were applied. The contact impedance of the electrodes was monitored during electrical stimulation for safety considerations. The *in vivo* measurements were conducted the day before the epilepsy resection surgery. The entire process took no more than half an hour, and the research protocol was integrated with clinical care. A pre-stimulation clinical assessment was performed by a physician before conducting the experiment. When conducting the experiment, physicians were present at the bedside during the entire procedure to monitor the subject’s clinical safety.

### 2.3 Intracranial recording protocol

The tES-induced intracranial voltage changes were measured usmg clinically implanted sEEG electrodes (Sinovation, China; 0.8 mm diameter, 3.5 mm contact spacing, 2 mm contact length), and the sites of electrode insertion were determined by clinical considerations. A clinical amplifier (EEG-1200C, Nihon Kohden, Japan) was used to collect multichannel tACS-induced voltages with band-pass filtered from 0.16 to 300 Hz and a sampling rate of 2000 Hz (AC recording mode). To record the tDCS induced voltage, some sEEG electrodes (less than 64 channels) were selected to be connected to an EEG amplifier (SynAmps2 RT, Neuroscan, USA), and the amplifier was set to DC - 200 Hz bandwidth and 2000 Hz sampling rate (DC recording mode). The ground electrode was positioned in the right mastoid region.

The tACS-induced voltage magnitudes were measured by fitting a sinusoid to the recordings during sinusoidal stimulation. We estimated the electric field component along the direction of the sEEG electrodes (projection of the electric field). The projected electric field was calculated in the direction of adjacent recording channels by subtracting the voltage values and dividing by their distance. For the induced voltages from the DC recording mode (SynAmps2 RT, Neuroscan), the DC bias and DC drift were observed because band-pass filtering was not used. The recording data were processed with demeaned (by subtracting the average voltage value) and detrended (by correcting for the slope of voltage) on each measuring channel. Because of the noisy interference, some recording channels were discarded.

### 2.4 Simulation model and data analysis

Individual pre-implantation Tl-weighted MRI images (TR= 8.2 ms, TE = 3.2 ms, flip angle =12°) and post-implantation CT images (237 mA/slice, 120 kV) were acquired using a 3T MRI scanner and CT scanner with a resolution of 0.5 × 0.5× 1.0 mm^3^. As depicted in Fig 1, the MRI and CT images were co-registered to determine the electrode insertion position usmg Brainstorm (brainstorm3, https://neuroimage.usc.edu/brainstorm/) [24]. The tES modeling process was performed using the open-source software package ROAST (ROAST 3.2, https://www.parralab.org/roast/) [25]. ROAST, called SPM12, to segment the MRI into grey matter (GM), white matter (WM), cerebrospinal fluid (CSF), bone, scalp, and air cavities. The conductivities of all tissues were assumed to be isotropic (WM: 0.126 S/m, GM: 0.276 S/m, CSF: 1.65 S/m, skull: 0.01 S/m, scalp: 0.465 S/m, air cavities: 2.5×10-14 S/m, electrode: 5.9×10^7^ S/m, gel: 0.3 S/m). Owing to the advantage of the minimally invasive method for sEEG, the sEEG electrodes were not modeled in our simulation model. The Pearson correlation coefficient was used for correlation analyses between different measured or predicted values. A best-fit line was obtained by linear regression for these values.

**Fig. 1.**
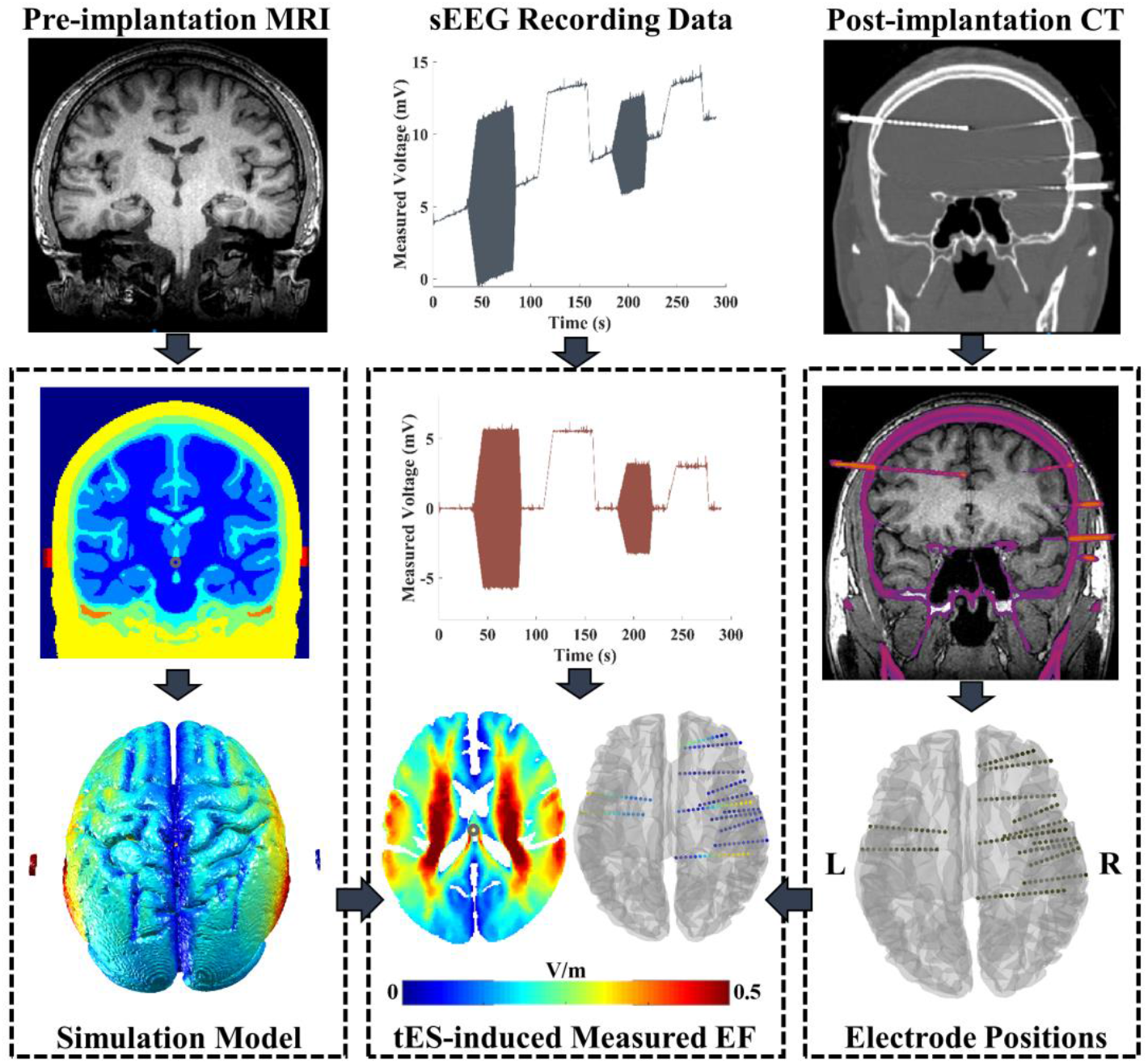
Electric field modeling and *in vivo* data analysis procedure. Brainstorm software was applied for individual post-implantation CT/pre-implantation MRI image registration. After manually editing the sEEG electrode site on the original CT, the positions of the electrode were exported in the MNI coordinates. ROAST was adopted to perform EF modeling by using individual MRI image. The voxel coordinates of the stimulation electrode were imported to ROAST, and then the stimulation electrodes were modeled at the customized locations. Combining with the simulated results and MNI coordinates of sEEG electrode, the voltage values of each electrode site were obtained and further compared with the measured value.

## 3. Results

Twelve epilepsy subjects were recruited, and the details of intracranial recording and stimulation montage are listed in Table I. Aa a result, multichannel tACS was applied to nine epilepsy subjects (S1-S9), and intracranial voltage changes were recorded with 964 sEEG electrodes in total. As shown in Supplementary Fig. I, these electrodes covered the subcortical brain area, extensive portions of the lateral and medial frontal, parietal and temporal cortex of the left and/or right hemisphere. The tDCS-induced voltage change was recorded with 706 sEEG electrodes in all subjects (S1-S12). Meanwhile, multichannel tDCS and tACS were applied to five subjects (S2, S5, S7, S8, and S9) under DC recording mode.

**Table I.** Summary of recording channels and stimulation montage for participants.

| Participants | Number of Recording Electrode |  | Position of Stimulation Electrode (10-10 EEG cap) |
| --- | --- | --- | --- |
|  | EEG-1200C | SynAmps2 RT |  |
| S1 | 91 | 61 | Electrode 1: T8, Electrode 2: Oz, Electrode 3: T7 |
| S2 | 108 | 58 | Electrode 1: T8, Electrode 2: Oz, Electrode 3: C5 |
| S3 | 173 | 62 | Electrode 1: T8, Electrode 2: Fpz, Electrode 3: T7 |
| S4 | 140 | 62 | Electrode 1: T8, Electrode 2: Fpz, Electrode 3: T7 |
| S5 | 46 | 46 | Electrode 1: T8, Electrode 2: Fpz, Electrode 3: T7 |
| S6 | 93 | 60 | Electrode 1: Fpz, Electrode 2: C3, Electrode 3: T8 |
| S7 | 117 | 64 | Electrode 1: T8, Electrode 2: C4, Electrode 3: Fp1, Electrode 4: T7 |
| S8 | 101 | 63 | Electrode 1: T7, Electrode 2: Fp1, Electrode 3: T8, Electrode 4: Oz |
| S9 | 95 | 56 | Electrode 1: T7, Electrode 2: C3, Electrode 3: T8 |
| S10 | Not used | 57 | Electrode 1: T7, Electrode 2: Fp2 |
| S11 | Not used | 60 | Electrode 1: T7, Electrode 2: T8 |
| S12 | Not used | 57 | Electrode 1: T7, Electrode 2: T8 |

### 3.1 Spatial linearity of intracranial voltages for multi-electrode stimulation

Linear superposition indicated that the multichannel tES-induced electric fields reflect the independent contribution from each bipolar montage. We verified this assumption using *in vivo* intracranial measurements. As shown in Fig. 2A, a four-electrode configuration was adopted for subject S7. When “Montage-14” and “Montage-23” were applied independently for S7, the total induced voltage showed a high correlation with the values from “Montage-13” and “Montage-24” (the Pearson correlation coefficient *r* = 0.99, *p* < 0.001; the slope of the linear fits= 1.00, Fig. 2B). Furthermore, the induced voltage by “Montage-3-12” was highly correlated with the sum of the voltage generated independently by the “Montage-31” and “Montage-32” for S7 *(r* = 0.99, *p* < 0.001; *s* = 0.99, Fig. 2C). Similar results were observed for subjects S1-S7 *(r* = 0.99, *p* < 0.001; *s* = 1.01, Fig. 2D). With *in vivo* measurements of tDCS induced voltage, we further found that, regardless of tDCS or tACS, the voltage induced by multichannel stimulation was highly correlated with the sum of the voltages generated independently by bipolar stimulation for S2, S5, S7-S9 (tDCS: *r* = 0.99, *p* < 0.001; tACS: *r* = *0*.*99,p* < 0.001; Fig. 2E-F).

**Fig. 2.**
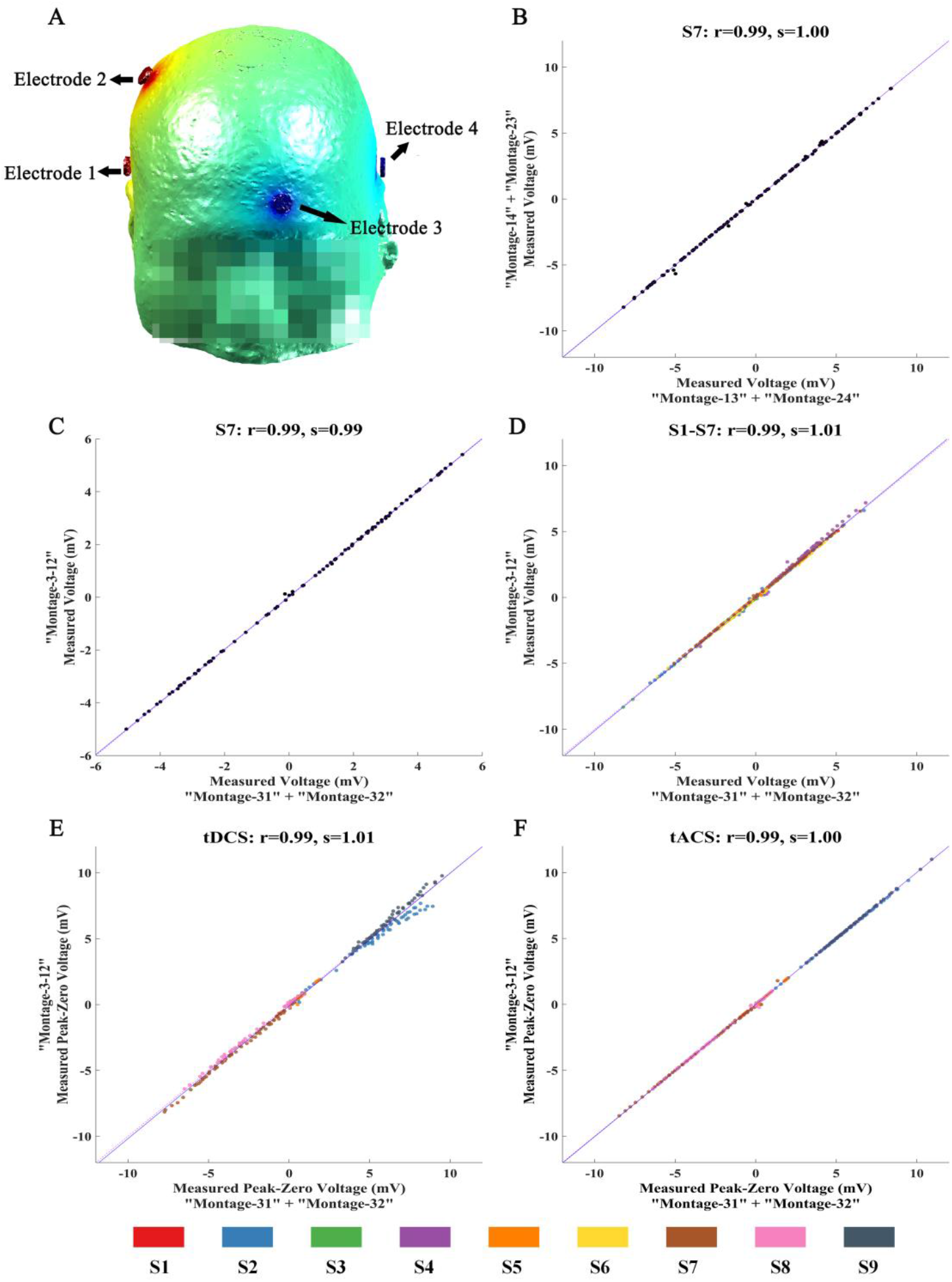
Spatial linearity of induced voltages for different electrode montage. **(A)** Placement position of stimulation electrode for S7. **(B-C)** Comparison of multichannel tES-induced voltages and the sum of the voltages generated independently by bipolar tES for S7 (AC recording mode). “Montage-31” represents electrode montage: Electrode3 as stimulation electrode (0.5 mA), Electrode! set as return electrode; “Montage-32”: Electrode3 as stimulation electrode (0.5 mA), Electrode2 set as return electrode; “Montage-3-12”: Electrode3 as stimulation electrode (1 mA), Electrode! set as return electrode (0.5 mA), Electrode2 set as return electrode (0.5 mA); **(D)** Comparison of induced voltages in all subjects (AC recording mode). **(E-F)** Comparison of multichannel tES-induced voltages and the sum of voltage generated independently by bipolar tES for S2, S5, S7-S9 (DC recording mode). The blue line represents fitting line. Points falling on the magenta line represent perfect prediction (slopes= 1).

### 3.2 Measurements and predictions of the electric field

To validate whether the computational model can accurately predict the electric field induced by multichannel tES, we compared the differences between the measured and simulated fields. The recorded voltages were highly correlated with the simulated voltage values for bipolar tES (Pearson correlation coefficient *r* = 0.90, *p* < 0.001; the slope of the best linear fit *s* = 0.93, Fig. 3A). Similar results were observed for multichannel tES *(r* = 0.92, *p* < 0.001; *s* = 0.97, Fig. 3C). A moderate correlation was also found between the measured and simulated EF for bipolar tES *(r* = *0*.*69,p* < 0.001; *s* = 0.74, Fig. 3B) and multichannel tES *(r* = 0.74, *p* < 0.001; *s* = 0.87, Fig. 3D). Furthermore, the correlation between the measured projected EFs and the simulated values was lower than that of the voltage value, which is consistent with a previous studies [17]. The correlation coefficient and slope of the linear fit between the measured and simulated values for each subject (S1-S9) are illustrated in Table II (for voltages: *r* = 0.92 ± 0.05, *s* = 0.95 ± 0.18; for projected electric fields: *r* = 0.71 ± 0.11, *s* = 0.81 ± 0.17).

**Table II.** The correlation coefficients and linear fit-slope during multichannel tES.

| Participants | Voltage |  | Projected EFs |  |
| --- | --- | --- | --- | --- |
|  | *r* | *s* | *r* | *s* |
| S1 | 0.94 | 0.83 | 0.61 | 0.75 |
| S2 | 0.96 | 0.94 | 0.63 | 0.68 |
| S3 | 0.90 | 1.21 | 0.90 | 0.87 |
| S4 | 0.94 | 1.26 | 0.64 | 1.21 |
| S5 | 0.88 | 0.75 | 0.56 | 0.69 |
| S6 | 0.97 | 0.77 | 0.75 | 0.68 |
| S7 | 0.97 | 0.97 | 0.76 | 0.95 |
| S8 | 0.91 | 0.98 | 0.81 | 0.78 |
| S9 | 0.80 | 0.87 | 0.76 | 0.72 |

*r*, *s* represent correlation coefficients and slope of a linear fit between measured and simulated values, respectively.

**Fig. 3.**
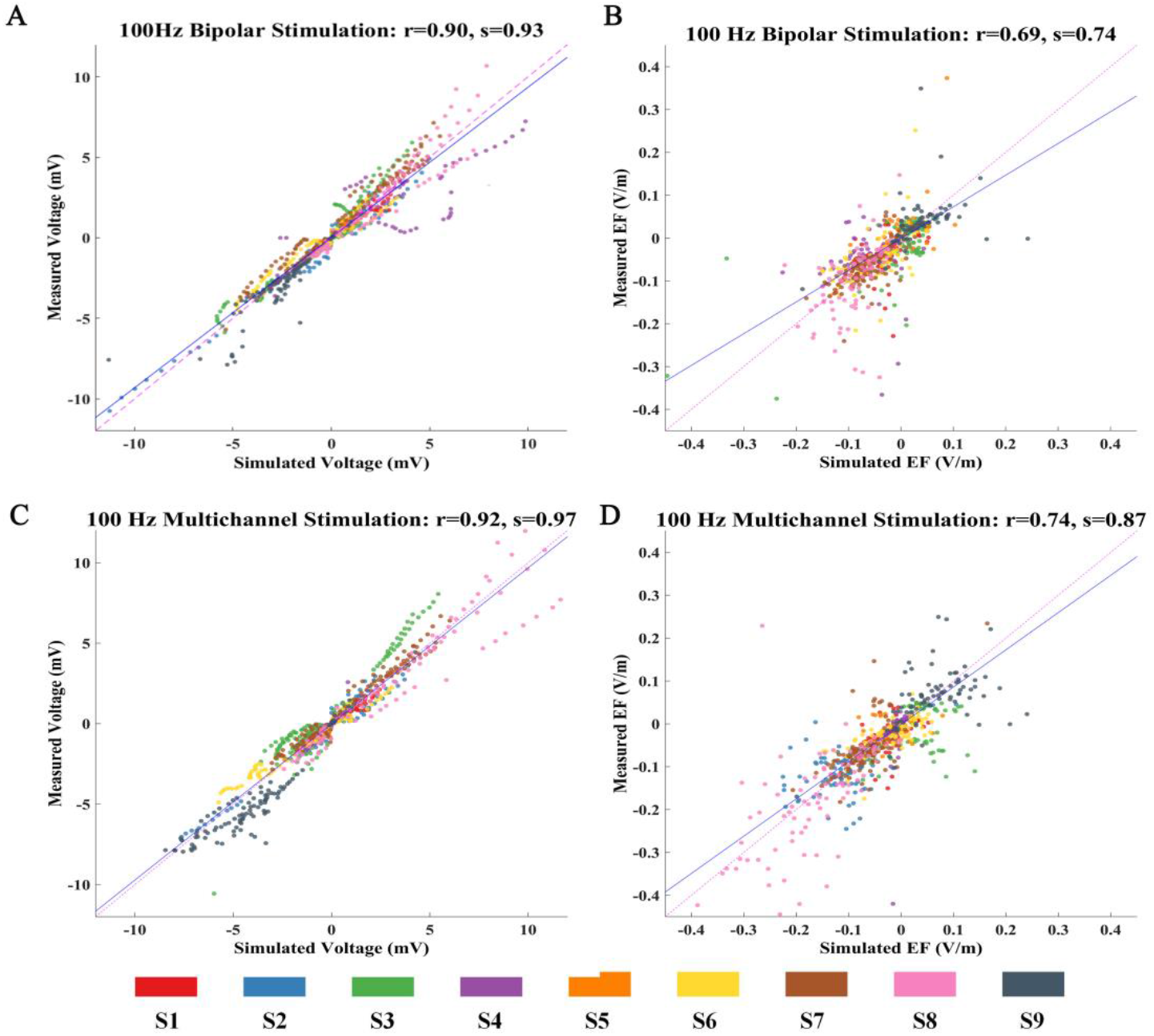
Correlation between simulated and measured values for 100 Hz stimulation. **(A-B)** Measured voltages and projected electric fields with simulated values from individualized model for bipolar stimulation. **(C-D)** Comparison of recorded voltages and projected electric fields with predicted values for multichannel stimulation.

### 3.3 Intracranial recording for tDCS and tACS

To compare the difference in induced EFs between tDCS and tACS, the same intensity tDCS and tACS were applied to all subjects, as displayed in Fig. 4A. With the equal-intensity stimulation, similar peak-zero voltage magnitudes were generated by tDCS and tACS within the brain, and tDCS-induced peak-zero voltages were highly correlated with the values of tACS *(r* = 0.99, *p* < 0.001; *s* = 0.99, Fig. 4B). On further comparing the measured voltage in different brain tissues, we observed that the voltage induced by equal-intensity tDCS and tACS had a high correlation within GM *(r* = 0.99, *p* < 0.001; *s* = 1.00, Fig. 4C) and WM *(r* = 0.99, *p* < 0.001; *s* = 1.02, Fig. 4D). Even when the polarity of tDCS was altered, the measured peak-zero voltage between these two tDCS conditions demonstrated a high correlation *(r* = *0*.*95,p* < 0.001; *s* = −0.98, Fig. 4E). We also calculated the projected EFs and found that the tDCS-induced EFs were moderately correlated with the values obtained by tACS *(r* = 0.79, *p* < 0.001; *s* = 0.78, Fig. 4F).

**Fig. 4.**
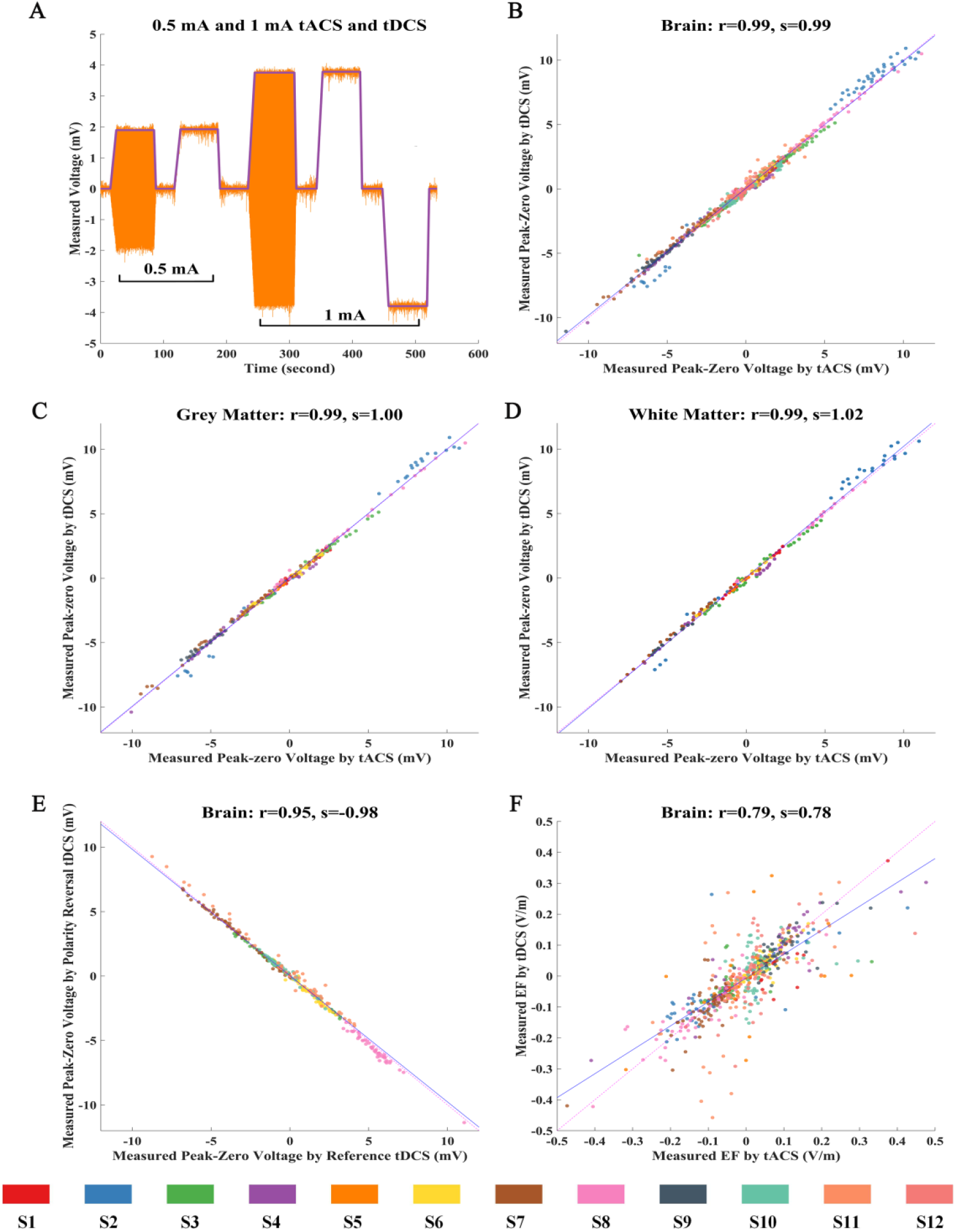
Comparison of tDCS-induced and tACS-induced field. **(A)** Measured voltages for different intensity tDCS and tACS. **(B)** Comparison of lmA tDCS-induced and tACS-induced voltage within the brain. **(C-D)** Comparison of tDCS-induced and tACS induced voltage within GM and WM. **(E)** Comparison of tDCS-induced voltage with opposite stimulation polarity. (F) Comparison of tDCS-induced and tACS-induced EF within the brain.

## 4. Discussion

In this study, we investigated multichannel tES-generated EF distributions by *in vivo* recordings. We found that the tES-induced voltage accords with the linear superposition within the human brain, which means that the multichannel tES-induced field is a summation of values induced by each independent bipolar stimulation. A previous study also supported this assumption at the scalp level using electroencephalography (EEG) [26]. Combined with our intracranial *in vivo* evidence, we can infer that linear superposition is tenable within intracranial and extracranial brain tissues. Compared with a previous multichannel tACS study in non-human primates [19], our study provides direct evidence for the underlying biophysical characteristics of multichannel tDCS and tACS in human brains. This is of great significance to tES optimization and clinical research.

In addition, by combining computational simulations based on individual MRI, we observed that the measured electric fields are highly correlated with the predicted ones in both bipolar and multi-electrode tES, which validates that the computational model is accurate to a certain extent in predicting the electric field distribution. Our results are consistent with those of previous *in vivo* studies [17]. However, individual variance still exists in the prediction accuracy of the simulation model. We infer that this may be related to the tissue conductivity differences, age differences, or electrode displacements. However, because of the limited number of subjects in the study, it was difficult to study the problem systematically. Notably, only a set of classic tissue conductivities was used in our simulation model, and the individual conductivity could be optimized to better match the actual recordings [27].

Furthermore, tDCS and tACS demonstrated almost the same induced peak-zero voltage magnitudes with the same stimulation intensity. To the best of our knowledge, this is the first report of tDCS-induced EF measurements across sEEG electrodes in humans. For these two electrical stimulation methods, tDCS and tACS showed different application characteristics. One of the most remarkable features is that direct currents were adopted in tDCS, while alternating current of different frequencies were applied in tACS. Therefore, the induced EFs were significantly different between the tDCS and tACS. The induced EF of tDCS is stable, whereas that of tACS is time-varying. When alternating current is applied sinusoidally in tACS, the current direction flips back and forth by 180° each half-wave, and its corresponding induced EF also exhibits sinusoidal variation over time. Thus, the tACS-generated EF was commonly represented by the peak EF value in most related modeling studies. However, there is still a lack of actual *in vivo* measurements to compare the differences between tACS and tDCS. Our study provides direct evidence for the comparative analysis of tACS and tDCS. In addition, tES modeling was calculated based on finite element analysis of quasi-static, and the induced field magnitudes were equal for tDCS and tACS at the same stimulation intensity. Our study is the first to support this hypothesis through experimental validation. Meanwhile, some previous studies also reported that the magnitudes of the measured voltages increased linearly with the stimulation intensity and changed only slightly when the tES stimulation frequency increased from 1 to 300 Hz [17, 21]. By analyzing these results, we can speculate that the head model can be considered as a purely resistive conductor and linear system during low-frequency electrical stimulation (0-200 Hz), where the propagative, inductive, and capacitive effects are negligible.

Notably, different electrical conductivities of brain tissues have been adopted for tDCS and tACS in some modeling studies [28, 29]. Until now, as a key factor for current flow modeling, most of the conductivity values in the literature were measured using alternating current, and the direct current conductivity are very lacking [30]. Our study found that tDCS and tACS induced similar peak-zero voltage magnitudes in the grey matter and white matter. Thus, the same tissue conductivity values can be adopted for tDCS or tACS modeling. In addition, we did not use any filtering to ensure that the DC voltage changes could be recorded, which led to a DC drift and bias for sEEG recordings. The noise of the recorded signal also increased. Thus, the correlation between tDCS and tACS-induced EF was lower than that of the voltage value. Thus, from our findings, tACS is a better choice for *in vivo* measurements and model validation.

Because of the limitations of the relevant conditions, this study has certain limitations. The sample size was small, and only twelve subjects were involved, which restrained the generalization of our results. Furthermore, our simulation head model only segments into common brain tissue components, a more detailed model that includes more subdivided tissue components would be desirable. Meanwhile, the conductivities of tissues employed in our study are widely used, and their accuracy can be further improved. Moreover, we did not use the isotropic simulation model in our study because of the lack of diffusion tensor imaging (DTI) data, so it is hard to evaluate the benefits of anisotropic modeling.

## 5. Conclusion

Our *in vivo* measurements indicate that the tES-induced electric field accords with the linear superposition and quasi-static assumption within the human brain. Meanwhile, we found that the simulation model of tES could reliably provide predicted values for bipolar and multi-electrode stimulation. Our study provides a solid foundation for tES modeling and clinical applications.

## Data availability

Data involved in this study are available upon reasonable request.

## Acknowledgements

We want to thank all subjects and their family members for their generous support and participation in this study.

